# Gut bacterial Infection drives Parkinsonian pathology in LRRK2 G2019S Knock-in Mice

**DOI:** 10.64898/2026.06.08.728789

**Authors:** Yuhang Wang, Matthew A. Weber, Nick C Stodden, Lily M Ziebarth, Sivakumar Lingappa, Monserrath A Borjabustamante, Tharoon A Balakrishnan, Manab Jaily, Taylor Sakaguchi, Ramasamy Thangavel, Wen Wen, Jia Luo, Nandakumar S. Narayanan, Zizhen Kang

**Author notes:** Corresponding author: Lead contact: Zizhen Kang, Nandakumar S. Narayanan.

## Abstract

The LRRK2 G2019S mutation is one of the most common genetic risk factors for Parkinson’s disease (PD), yet LRRK2 G2019S knock-in (KI) mice rarely develop robust neurodegeneration under basal conditions, suggesting that additional environmental triggers are required for disease progression. Here, we established a clinically relevant gene–environment interaction mouse model of PD by subjecting LRRK2 G2019S KI mice to recurrent *Citrobacter (C.) rodentium* infection, a murine model of enteric bacterial inflammation. Repeated infection induced progressive PD-like phenotypes selectively in KI mice, including motor impairment, reduced locomotor activity, impaired motor coordination, selective nigrostriatal dopaminergic neurodegeneration, enhanced neuroinflammation, and pathological phosphorylated α-synuclein (p-αSyn) accumulation, whereas wild-type (WT) mice remained largely resistant. Mechanistically, infected KI mice developed markedly exacerbated colonic inflammation, epithelial barrier dysfunction, increased intestinal permeability, and enhanced inflammasome activation despite normal bacterial clearance, indicating that pathogenic LRRK2 signaling amplifies inflammatory responses rather than impairing antimicrobial defense. In parallel, recurrent infection induced pronounced intestinal p-αSyn accumulation and expansion of pathology beyond the epithelial layer in KI mice, supporting a gut–brain axis mechanism linking intestinal inflammation to neurodegeneration. Collectively, these findings demonstrate that the LRRK2 G2019S mutation functions as a sensitizing factor that cooperates with recurrent enteric inflammation to drive PD-related pathology. This study establishes a physiologically relevant LRRK2 G2019S gene– environment interaction mouse model that recapitulates key behavioral, neuropathological, and inflammatory features of PD.

## Introduction

Parkinson’s disease (PD) is the second most common neurodegenerative disorder worldwide and is characterized by progressive degeneration of dopaminergic neurons in the nigrostriatal pathway, leading to the cardinal motor symptoms of bradykinesia, rigidity, resting tremor, and postural instability. More than one million individuals in the United States are currently affected by PD, and its global prevalence more than doubled between 1990 and 2016, exceeding six million cases worldwide(1, 2). With rapidly aging populations, the global burden of PD is projected to increase substantially in the coming decades. Neuropathologically, PD is defined by the presence of Lewy bodies and Lewy neurites, intracellular proteinaceous inclusions enriched in misfolded and aggregated α-synuclein (α-syn). Among the various pathological forms of α-syn, phosphorylation at serine 129 (pS129) represents the predominant species within Lewy pathology and is widely regarded as a hallmark of disease-associated α-syn aggregation and neurodegeneration.

Genetic susceptibility contributes substantially to PD risk, with current estimates suggesting that up to 15% of patients harbor pathogenic genetic variants(3). Among these, mutations in the leucine-rich repeat kinase 2 (LRRK2) gene represent the most common cause of monogenic PD. The LRRK2 G2019S mutation, located within the kinase domain, is detected in approximately 1– 3% of sporadic PD cases and 4–8% of familial cases(4, 5). Consequently, targeting LRRK2 kinase activity has become an area of intense therapeutic interest. Despite its strong association with human disease, LRRK2 G2019S knock-in (KI) mice typically develop only subtle or prodromal PD-like phenotypes, suggesting that the mutation alone is insufficient to drive full disease manifestation and that additional environmental or inflammatory triggers may be required to unmask pathogenic neurodegeneration(6). Notably, LRRK2 is broadly expressed in peripheral tissues, with particularly high expression in immune cells, kidneys, and lungs, and lower but functionally significant expression in the brain. Accumulating evidence implicates LRRK2 in the regulation of immune signaling, endolysosomal trafficking, ciliogenesis, and protein translation, linking its diverse cellular functions to both central nervous system pathology and peripheral immune mechanisms relevant to PD(7).

The pathogenesis of PD remains incompletely understood. Intriguingly, many PD patients exhibit gastrointestinal dysfunction accompanied by intestinal inflammation and α-syn pathology within enteric neurons, abnormalities that often precede the onset of classical motor symptoms by several years(8–10). Growing evidence from both human studies and animal models supports a critical role for the gut–brain axis in PD initiation and progression(11, 12). This concept is exemplified by the Braak hypothesis, which proposes that α-syn pathology can originate in the gastrointestinal tract and spread in a stereotypical manner via the vagus nerve to the ventral midbrain, ultimately leading to the degeneration of dopaminergic neurons in the substantia nigra pars compacta(13). Consistent with this model, several population-based cohort studies have demonstrated that individuals with inflammatory bowel disease (IBD) have a significantly elevated risk of developing PD, with increases ranging from 20% to 90% compared with non-IBD populations(14). Importantly, anti-inflammatory treatment of IBD has been associated with a reduced risk of subsequent PD development(15).

Despite these observations, the molecular pathways linking intestinal inflammation to neurodegeneration remain poorly defined. Notably, genome-wide association studies have identified strong genetic associations between LRRK2 and IBD(16). Furthermore, recent exome-sequencing analyses have uncovered functional LRRK2 variants shared between IBD and PD, suggesting that LRRK2 may function as a signaling hub linking intestinal inflammation to neurodegenerative processes(17). Consistent with this possibility, we and others have shown that LRRK2 promotes intestinal inflammation in experimental colitis models(18, 19). In addition, recent studies demonstrated that gut inflammation induced by dextran sodium sulfate (DSS) promotes parkinsonian phenotypes in LRRK2 G2019S transgenic and knock-in mice(20, 21), which otherwise exhibit minimal neurological symptoms in the absence of inflammatory stimuli. These findings suggest that intestinal inflammation may act as an environmental modifier that increases the penetrance of parkinsonism in genetically susceptible hosts.

Intestinal infections represent a major cause of gut inflammation. Beyond causing acute illnesses such as fever and diarrhea, intestinal infections may also contribute to the development of chronic inflammatory and systemic diseases, including IBD, metabolic disorders, and neurodegenerative diseases such as PD. Moreover, PD patients frequently exhibit gut microbiota dysbiosis(14), and transplantation of microbiota from PD patients promotes parkinsonian pathology in germ-free α- synuclein-overexpressing mice(22). However, whether and how intestinal infection promotes LRRK2-associated parkinsonism remains largely unknown.

*Escherichia coli*, a member of the Enterobacteriaceae family, is a common enteric pathogen capable of triggering intestinal infection and inflammation. Notably, Enterobacteriaceae species are frequently enriched in the gut microbiota of PD patients(23). *Citrobacter rodentium (C. rodentium)* is a natural murine pathogen that closely models human enteropathogenic *E. coli* infection, making it a highly relevant system for studying host–pathogen interactions at the intestinal barrier and their systemic consequences(24).

We therefore hypothesized that intestinal infection with *C. rodentium* promotes parkinsonian pathology in LRRK2 G2019S KI mice. To test this hypothesis, LRRK2 G2019S KI mice were subjected to repeated cycles of *C. rodentium* infection. We found that recurrent infection induced progressive PD-like behavioral deficits, dopaminergic neurodegeneration, and pathological accumulation of phosphorylated α-syn. In parallel, KI mice exhibited enhanced late-phase intestinal p-αSyn aggregation, and heightened acute intestinal inflammation and inflammasome activation in the gut, following bacterial challenge. Together, these findings support a gut–brain axis mechanism whereby intestinal inflammation amplifies neurodegenerative pathology. Collectively, our results reveal a potent gene–environment–immune interaction driving PD pathogenesis and establish an infection-augmented LRRK2 G2019S mouse model for mechanistic and therapeutic studies of parkinsonism.

## Materials and Methods Animals

LRRK2 G2019S KI (LRRK2 KI) mice were purchased from Jackson Laboratory (JAX030961). LRRK2 KI strain is congenic on C57BL/6J background. To get littermate wild type control mice, LRRK2^KI/+^ male mice were mated with LRRK2^KI/+^ female mice, sex matched LRRK2^+/+^ mice (Referred to as WT control) and LRRK2^KI/KI^ mice (Referred to as LRRK2 KI) were used at the age of 8-12 weeks. Both male and female mice were used for the experiments. All mice were bred and maintained in individually ventilated cages under specific pathogen-free conditions in accredited animal facilities. Animal experiments were approved by the Institutional Animal Care and Use Committee of the University of Iowa.

### *C. rodentium* culture and mouse infection

*Citrobacter rodentium* (*C. rodentium*, ATCC 51459) was purchased from ATCC. Bacteria were recovered from frozen glycerol stocks and cultured overnight in Luria–Bertani (LB) broth at 37 °C with shaking. On the day of infection, the overnight culture was diluted 1:10 into fresh LB broth and expanded for approximately 4 h at 37 °C with shaking until reaching an OD600 of approximately 0.6–0.8. Bacteria were collected by centrifugation, washed twice with sterile PBS, and resuspended in PBS for oral gavage. Bacterial concentration was estimated by OD600 measurement and confirmed by serial dilution plating on MacConkey agar.

LRRK2 KI mice and WT littermate controls (8–12 weeks old) were fasted for 4 h prior to infection and orally gavaged with approximately 4 × 10⁸ colony-forming units (CFU) of *C. rodentium* in 200 μL PBS, while control animals received 200 μL PBS alone. Following infection, mice were monitored daily for body weight, stool consistency, and clinical signs of disease. Fecal pellets were collected at indicated time points, homogenized in sterile PBS, serially diluted, and plated on MacConkey agar for quantification of bacterial burden. To establish the PD-like mouse model, mice were subjected to four repeated cycles of infection at approximately one-month intervals, using a repeated intestinal infection paradigm adapted from a previously established infection-induced model of Parkinson’s disease-like pathology in *Pink1*-deficient mice (25). For acute infection experiments, mice were sacrificed at the indicated time points following a single infection as described in the figure legends.

## Intestinal permeability assay

Nine days after *C. rodentium* infection, intestinal barrier integrity was assessed using a fluorescein isothiocyanate–dextran (FITC–dextran) permeability assay (average molecular weight 4 kDa; Sigma-Aldrich, USA) as previously described(18). Briefly, mice were fasted for 4 h and then orally gavaged with FITC–dextran at a dose of 60 mg per 100 g body weight. Four hours after administration, mice were euthanized and blood was collected via cardiac puncture. Blood samples were kept at room temperature for 2 h protected from light, and serum was obtained by centrifugation at 10,000 × g for 10 min. Serum fluorescence intensity was measured using a SpectraMax i3 plate reader (Molecular Devices, San Jose, CA, USA). FITC–dextran concentrations were calculated based on a standard curve generated from serial dilutions of FITC–dextran.

## Colonic explant culture

Whole colons were harvested from LRRK2 WT or LRRK2 G2019S KI mice, thoroughly rinsed with serum-free DMEM medium, and weighed to determine their initial weight. The collected colon tissues were cut into 2 mm pieces and then cultured as explants in regular RPMI 1640 medium supplemented with 10% FBS, L-glutamine, penicillin, and streptomycin, and placed in a standard cell-culture incubator for 24 hours. After culture, cell-free supernatants were collected by centrifugation at 12,000 × g for 10 min at 4°C, aliquoted, and stored at −20°C. Cytokine levels were subsequently analyzed.

## Microdissection of Midbrain Dopaminergic Regions

Mice were euthanized and transcardially perfused with ice-cold phosphate-buffered saline (PBS). Brains were rapidly harvested and placed on an ice-cold dissection platform. Coronal sections encompassing the midbrain were prepared, and the substantia nigra pars compacta (SNpc) and ventral tegmental area (VTA) were identified based on anatomical landmarks from the mouse brain atlas (approximately bregma −2.8 to −3.8 mm). These regions were bilaterally microdissected under a stereomicroscope using fine forceps and microsurgical blades, immediately snap-frozen on dry ice, and stored at −80°C until protein and RNA analyses.

## Immunohistochemistry

On the day of sacrifice, mice were deeply anesthetized and transcardially perfused with cold phosphate-buffered saline (PBS) for 1 minute to remove intravascular blood. Brains were rapidly dissected and fixed in 4% paraformaldehyde (PFA) at 4 °C for 24 hours. Following fixation, tissues were cryoprotected by immersion in 30% sucrose in PBS at 4 °C until fully equilibrated, embedded in optimal cutting temperature (OCT) compound, and sectioned coronally at a thickness of 40 μm using a cryostat (Leica, CM1860). Free-floating sections were permeabilized and blocked in PBS containing 0.3 percent Triton X-100 and 5% normal goat serum at 37 °C for 1 hour to reduce nonspecific binding. Sections were then incubated overnight at 4 °C with primary antibodies (**Supplemental table 1**). After primary antibody incubation, sections were washed three times with PBS for 10 minutes each and incubated with species-appropriate secondary antibodies (**Supplemental table 2**) for 1 hour at room temperature in the dark. All antibodies were diluted 1:1000 in PBS before use. Sections were subsequently washed three additional times with PBS, counterstained with DAPI for 5 minutes to visualize nuclei, mounted onto glass slides, and coverslipped using an antifade mounting medium (Epredia™ Immu-Mount™, Cat#9990412).

Formalin-fixed, paraffin-embedded colon and distal small intestine tissues were sectioned at 5 μm thickness and processed for immunohistochemistry. Sections were deparaffinized in xylene (three changes, 5 minutes each), rehydrated through a graded ethanol series (100%, 95%, 70%, and 50%), and rinsed in distilled water. Antigen retrieval was performed by heating sections in citrate buffer (pH 6.0) at 95 °C for 30 minutes in a pressure cooker, followed by gradual cooling to room temperature. After antigen retrieval, sections were permeabilized with 0.3% Triton X-100 in blocking buffer (5% normal goat serum in PBS) for 1 hour at room temperature. Sections were then incubated overnight at 4 °C with primary antibody against phosphorylated α-synuclein (AB51253, Abcam) diluted 1:1000 in blocking buffer. The following day, sections were washed three times with PBS and incubated with appropriate fluorescence-conjugated secondary antibodies for 1 hour at room temperature in the dark. After additional PBS washes, nuclei were counterstained with DAPI, and sections were mounted using an antifade mounting medium (Immu-Mount, Epredia). All sections were imaged using VS-ASW-S6 imaging software (Olympus, Center Valley, PA) and an Olympus Slide Scanner (Olympus VS120) with a 20x objective.

## Quantification of TH, p-αSyn, IBA1, and GFAP immunoreactivity

Immunofluorescence images of the substantia nigra and striatum were acquired using identical microscope settings across all experimental groups, including exposure time, gain, illumination intensity, and objective magnification. Quantification of TH, p-αSyn, IBA1, and GFAP immunoreactivity was performed using Olympus cellSens software. For each image, regions of interest (ROIs) were manually delineated based on anatomical landmarks corresponding to the ventral tegmental area (VTA), substantia nigra pars compacta (SNpc), and striatum. Immunoreactivity for each marker was quantified as the mean fluorescence intensity per unit area (mm²) within the selected ROI.

### Colon histological analysis after *C. rodentium* infection

Nine days after *C. rodentium* infection, mice were euthanized, and the colon was harvested and rinsed with ice-cold PBS. A distal 0.5-cm segment of the colon was collected, fixed in 10% neutral-buffered formalin for at least 24 h, paraffin-embedded, sectioned at 5 μm thickness, and stained with hematoxylin and eosin (H&E). Histological sections were evaluated in a blinded manner using a semi-quantitative scoring system adapted from published criteria(26). Four parameters were assessed: (i) epithelial damage, including crypt hyperplasia and goblet cell depletion, scored as 0 (none), 1 (mild), 2 (moderate), and 3 (severe); (ii) inflammatory cell infiltration in the lamina propria, scored as 0 (none), 1 (mild leukocyte infiltration or increased lymphoid follicles), 2 (moderate infiltrate with crypt broadening), and 3 (dense inflammatory infiltrate); (iii) extent of tissue involvement, scored as 0 (none), 1 (0–25%), 2 (25–50%), and 3 (>50% of the section affected); and (iv) severe pathological features, including submucosal inflammation, crypt abscesses, crypt branching, and ulceration or fibrosis, graded from 0 to 3 based on severity and presence. Scores from all categories were summed to yield a total histological score ranging from 0 to 12 for each section.

### Quantification of *C. rodentium* burden

On days 7, 12, 21, and 28 post-infection (p.i.), fresh fecal samples were collected, diluted in 1 mL PBS, and homogenized by bead beating with 1 mm ceramic beads for 40 s at 6000 rpm using a Vortex mixer (*BioSpec*). To determine colony-forming units (CFUs), serial dilutions of the homogenized samples were plated on MacConkey agar plates and incubated overnight at 37°C. C. rodentium colonies were identified based on their characteristic morphology, and CFUs were calculated and normalized to the weight (mg) of each fecal or tissue sample.

## Behavioral test Open field test

Open field activity was performed to evaluate exploratory behavior and spontaneous locomotor activity at three time points, 2, 4, and 6 mpi, in accordance with previously published methods. On each testing day, mice were acclimated to the behavioral testing room for at least 30 minutes before the experiment. Each animal was then placed individually into the center of a square open field arena, and its movement was recorded for a defined testing period using an automated video tracking system. Total distance traveled, average velocity, and time spent in the center versus peripheral zones were quantified as measures of locomotion and anxiety-like behavior. The apparatus was cleaned with 70% between trials to eliminate olfactory cues.

## Rotarod

The rotarod test was performed to assess motor coordination and motor learning. The procedure consisted of a training phase followed by a test phase. During training, mice were placed on a rotating rod (Columbus Instruments) operating at a constant speed of 4 rpm and were trained until they were able to remain on the rod for at least 20 seconds. The test phase was conducted 24 hours after training. For testing, the rotarod was set to accelerate gradually from 4 to 40 rpm over 300 seconds. Mice were placed on the rod at an initial speed of 4 rpm, and the time each animal remained on the rod before falling was recorded as the latency to fall, expressed in seconds.

## Grip strength test

Forelimb grip strength was assessed using a grip strength meter (Bio-GS3, BioSeb, France). Each mouse underwent three consecutive trials, and the mean value was calculated for analysis. Body weight was measured prior to testing. Given the absence of significant differences among groups, grip strength values were not normalized to body weight.

## ELISA

Cytokine production in the supernatants of colon explant tissues from mice was quantified using ELISA kits (R&D Systems) according to the manufacturer’s instructions. The following kits were used for specific cytokines: IL-1β (DY401-050) and IL-18 (7625–05). Cytokine levels were normalized to the weight of the corresponding colon tissue samples.

## Western blot

Colon tissues or midbrain dopaminergic region tissues were homogenized in radioimmunoprecipitation assay (RIPA) buffer supplemented with Complete Mini Protease Inhibitor Cocktail and Phosphatase Inhibitor Cocktail (Roche). Lysates were incubated on ice for 30 min with vortexing every 5 min, followed by centrifugation at 13,000 rpm for 10 min at 4°C. The supernatants were collected, and protein concentrations were determined using a BCA Protein Assay Kit (Pierce). Equal amounts of protein (30-50 μg per sample) were loaded onto 12% SDS-PAGE gels together with 5 μL of a prestained protein molecular weight marker (F4005, APExBio; 1610374, Bio-Rad). Proteins were separated by SDS-PAGE and transferred onto 0.45-μm PVDF membranes. For immunoblot analysis, primary antibodies against GSDMD (ab209845, Abcam), tyrosine hydroxylase (TH; AB152, Sigma-Aldrich), and β-actin (sc-47778, Santa Cruz Biotechnology) were used at a dilution of 1:1000. Horseradish peroxidase (HRP)-conjugated secondary antibodies were applied according to the host species of the primary antibodies. Membranes were incubated with enhanced chemiluminescence (ECL) reagent (34580, Thermo Fisher Scientific), and signals were detected using either X-ray film (Fuji X-Ray Film, Z&Z Medical) or a Bio-Rad ChemiDoc MP imaging system.

## Quantitative PCR

Colon tissues or midbrain tissues were carefully collected and homogenized using TRIzol reagent (Invitrogen) to ensure efficient RNA extraction. RNA was isolated according to the manufacturer’s instruction. The high-quality RNA (500 ng) was then reverse transcribed into complementary DNA (cDNA) using the High-Capacity cDNA Reverse Transcription Kit (Applied Biosystems). Quantitative PCR (Q-PCR) was subsequently carried out using SYBR Green Real-time PCR Master Mix (K1070, APExBio) on a Real-Time PCR System (Applied Biosystems). Each reaction was performed in triplicate in a 10 μl volume, containing 5 μl of SYBR Green mix, 0.3 μM of forward and reverse primers, and 0.5 μl of cDNA. The relative gene expression levels were calculated using the comparative Ct (ΔΔCt) method, normalized to the expression of a housekeeping gene β-actin.

## Statistics

Data were analyzed using two-way ANOVA followed by Bonferroni’s post hoc test. Statistical analyses were performed using GraphPad Prism (GraphPad Software, Inc., San Diego, CA). Data are presented as mean ± standard error of the mean (SEM). A *p* value ≤ 0.05 was considered statistically significant.

## Results

### *C. rodentium* infection induces motor impairment in LRRK2 KI mice

The LRRK2 G2019S mutation is one of the most common genetic risk factors for Parkinson’s disease (PD). However, knock-in (KI) mouse models carrying this mutation typically do not develop robust PD-like motor phenotypes under basal conditions, suggesting that genetic susceptibility alone is insufficient to drive overt disease. Environmental factors are increasingly recognized as critical contributors to PD pathogenesis, particularly in genetically predisposed individuals. Therefore, we hypothesized that additional environmental triggers are required to induce disease-relevant pathology in LRRK2 G2019S KI mice. Intestinal infection, a potent inducer of sustained gut inflammation, has emerged as a potential environmental driver of PD-related neurodegeneration through the gut–brain axis. Accordingly, we proposed that recurrent intestinal infection may act as a trigger to unmask latent vulnerability in LRRK2 G2019S KI mice.

To test this, we subjected LRRK2 KI mice and WT littermates to four repeated cycles of *C. rodentium* infection, with PBS-treated cohorts serving as controls, and longitudinally assessed behavioral outcomes (**Fig. 1A**). Behavioral testing was performed at 2-, 4-, and 6-months post-infection (mpi) to capture the temporal evolution of disease-relevant phenotypes.

**Figure 1.**
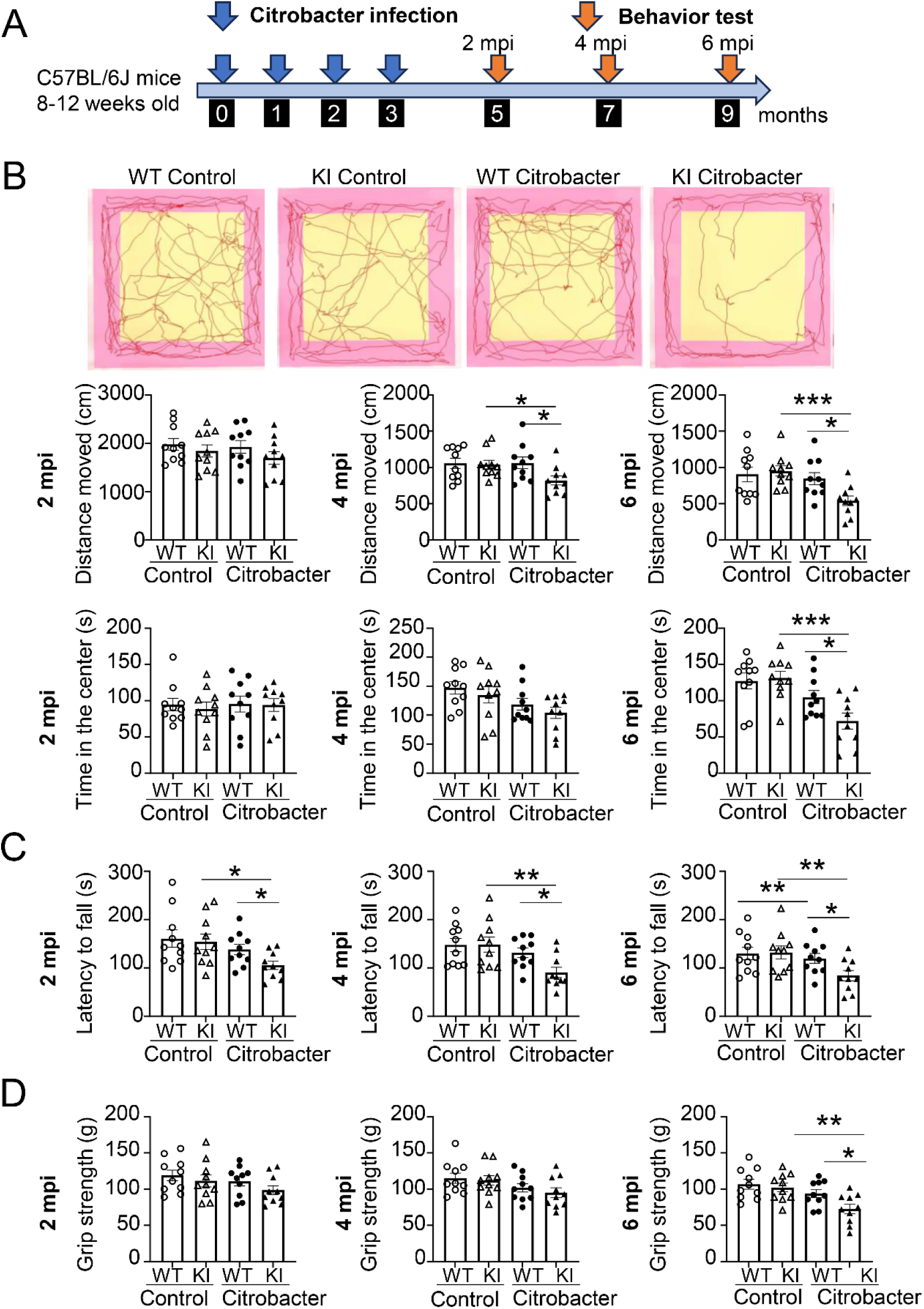
*C. rodentium* infection promotes motor impairment in LRRK2 G2019S KI mice. (A) Experimental design. Eight- to twelve-week-old LRRK2 wild-type (WT) and LRRK2 G2019S knock-in (KI) C57BL/6J mice were orally gavaged with 4 × 10⁸ CFU of *C. rodentium* per cycle for four consecutive infection cycles, with one-month intervals between cycles. Behavioral assessments were conducted at 2-, 4-, and 6-months post-infection (mpi). Age- and genotype- matched uninfected littermates served as controls (n = 10 per group). (B) Open field test. Representative movement traces at 6 mpi (top), with quantification of total distance traveled (middle) and time spent in the center area (bottom) at 2, 4, and 6 mpi. (C) Rotarod test. Motor coordination and balance were assessed by latency to fall (seconds) at indicated time points. (D) Grip strength test. Forelimb grip strength was measured and expressed as gram-force (g). Data are shown as mean ± SEM. *p<0.05, **p<0.01 and ***p<0.001. Data are representative of three independent experiments.

Spontaneous locomotor activity and exploratory behavior were first evaluated using the open-field assay. At 2 mpi, no significant differences were observed between groups. However, at 4 and 6 mpi, infected KI mice exhibited a marked reduction in total distance traveled compared with uninfected KI controls (**Fig. 1B**), indicating a progressive decline in locomotor activity. In parallel, infected KI mice spent significantly less time in the center zone at these later time points (**Fig. 1B**), suggesting reduced exploratory behavior, potentially reflecting increased anxiety-like behavior and/or impaired motor capacity.

Deficits in motor coordination emerged earlier than changes in locomotion. In the rotarod assay, infected KI mice displayed a significantly reduced latency to fall compared with uninfected KI controls as early as 2 mpi, with this impairment persisting through 4 and 6 mpi (**Fig. 1C**). These findings indicate an early and sustained disruption of motor coordination, balance, and motor learning. In contrast, reductions in muscular strength developed later. Forelimb grip strength was comparable between groups at 2 and 4 mpi but was significantly decreased in infected KI mice at 6 mpi (**Fig. 1D**), consistent with progressive neuromuscular dysfunction. Importantly, *C. rodentium* infection did not induce detectable behavioral abnormalities in WT mice, and no differences were observed between WT and KI mice under control conditions across all assays (**Fig. 1B-D**).

Collectively, progressive behavioral impairments were restricted to infected KI mice, which supports our hypothesis that a gene–environment interaction model in which recurrent intestinal infection functions as a triggering factor that unmasks latent neurodegenerative vulnerability in LRRK2 G2019S KI mice.

## Intestinal infection induces dopaminergic neurodegeneration in LRRK2 KI mice

Given the progressive motor deficits observed in LRRK2 G2019S KI mice at 6 months post-infection (mpi), we next tested whether these behavioral abnormalities were associated with degeneration of the nigrostriatal dopaminergic system. To address this, we assessed tyrosine hydroxylase (TH), the rate-limiting enzyme in dopamine synthesis and a canonical marker of dopaminergic neurons, by immunohistochemistry.

TH immunoreactivity was examined in midbrain regions, with a focus on the substantia nigra pars compacta (SNpc), the primary site of dopaminergic neuron degeneration in PD and the striatum, the major projection target of SNpc neurons. We also assessed TH immunoreactivity in the ventral tegmental area (VTA), another region of the midbrain that contains dopaminergic cell bodies. Strikingly, infected KI mice exhibited a marked loss of TH-positive neuronal cell bodies in the SNpc compared with both infected WT mice and uninfected KI controls (**Fig. 2A**). Quantitative analysis confirmed a significant reduction in TH signal intensity in the SNpc (**Fig. 2B**). In contrast, TH immunoreactivity in the VTA remained unchanged (**Fig. 2B**), indicating region-specific vulnerability of nigral dopaminergic neurons.

**Figure 2.**
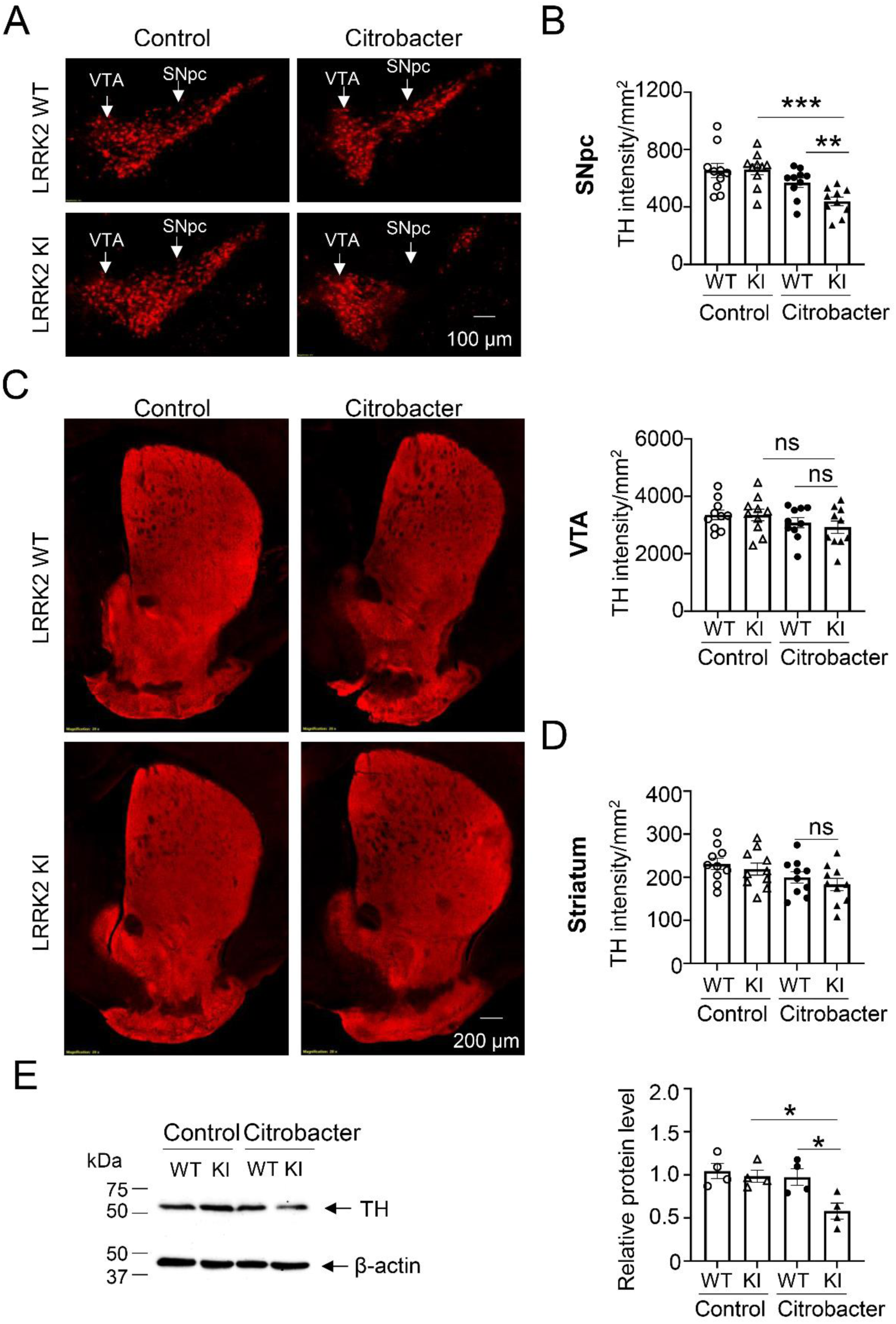
*C. rodentium* infection promotes dopaminergic neurodegeneration in LRRK2 G2019S KI mice. Tyrosine hydroxylase (TH) expression was assessed in the substantia nigra pars compacta (SNpc), ventral tegmental area (VTA), and striatum of LRRK2 WT and LRRK2 G2019S KI mice at 6 months post-infection (mpi). Age- and genotype-matched uninfected littermates served as controls. (A) Representative immunofluorescence images of TH-positive neurons in the SNpc and VTA from WT and KI mice under control and *C. rodentium* infection conditions. Scale bar, 100 μm. (B) Quantification of TH immunoreactivity in the SNpc (upper) and VTA (Bottom), expressed as fluorescence intensity per mm^2^. (C) Representative images of TH staining in the striatum. Scale bar, 200 μm. (D) Quantification of TH immunoreactivity in the striatum, expressed as fluorescence intensity per mm^2^. (E) Western blot analysis of TH protein levels in the midbrain dopaminergic regions, with β-actin as an internal control (left), and corresponding quantification using Image J (right). B and D, n = 10 mice per group for imaging analyses; E, n = 4 per group for western blot. Data are shown as mean ± SEM. *p<0.05, **p<0.01 and ***p<0.001. Data are representative of three independent experiments.

We next assessed dopaminergic projections in the striatum. Despite the pronounced loss of TH-positive neurons in the SNpc, TH-positive fibers in the striatum were largely preserved, with no significant reduction in signal intensity detected (**Fig. 2C, D**), although a modest decreasing trend was observed. These findings suggest that striatal dopaminergic terminals may be relatively preserved at this stage of disease progression compared with changes observed in the SNpc.

To independently validate these observations, TH protein levels in the midbrain were measured by Western blot. Consistent with the histological data, infected KI mice exhibited a significant reduction in TH protein expression compared with both infected WT and uninfected KI mice (**Fig. 2E**). Importantly, no significant differences were observed between WT and KI mice under control conditions, and intestinal infection alone did not induce dopaminergic loss in WT mice (**Fig. 2**). Thus, neurodegeneration was restricted to infected KI mice, indicating that intestinal infection is not sufficient to drive dopaminergic degeneration in the absence of genetic susceptibility.

Collectively, these findings demonstrate that recurrent intestinal infection selectively induces degeneration of SNpc dopaminergic neurons in LRRK2 G2019S KI mice, providing a neuropathological basis for the observed motor deficits.

## Intestinal bacterial infection promotes neuroinflammation in LRRK2 KI mice

We next investigated whether intestinal infection elicits neuroinflammatory responses in brain regions relevant to nigrostriatal pathology. Microglia and astrocytes are the principal innate immune effectors in the central nervous system and play critical roles in mediating neuroinflammatory responses. To assess glial activation, we performed immunohistochemical staining for IBA1 and GFAP, markers of microglia and astrocytes, respectively.

Following *C. rodentium* infection, both WT and LRRK2 KI mice exhibited increased IBA1 and GFAP immunoreactivity compared with their respective control groups, indicating that intestinal infection alone is sufficient to induce neuroinflammation (**Fig. 3B, 3D**). Notably, this response was significantly amplified in LRRK2 KI mice, which showed higher IBA1-positive microglial activation and GFAP-positive astrocytic reactivity compared with infected WT mice (**Fig. 3A–D**), indicating that pathogenic LRRK2 signaling enhances the neuroinflammatory response to intestinal infection.

**Figure 3.**
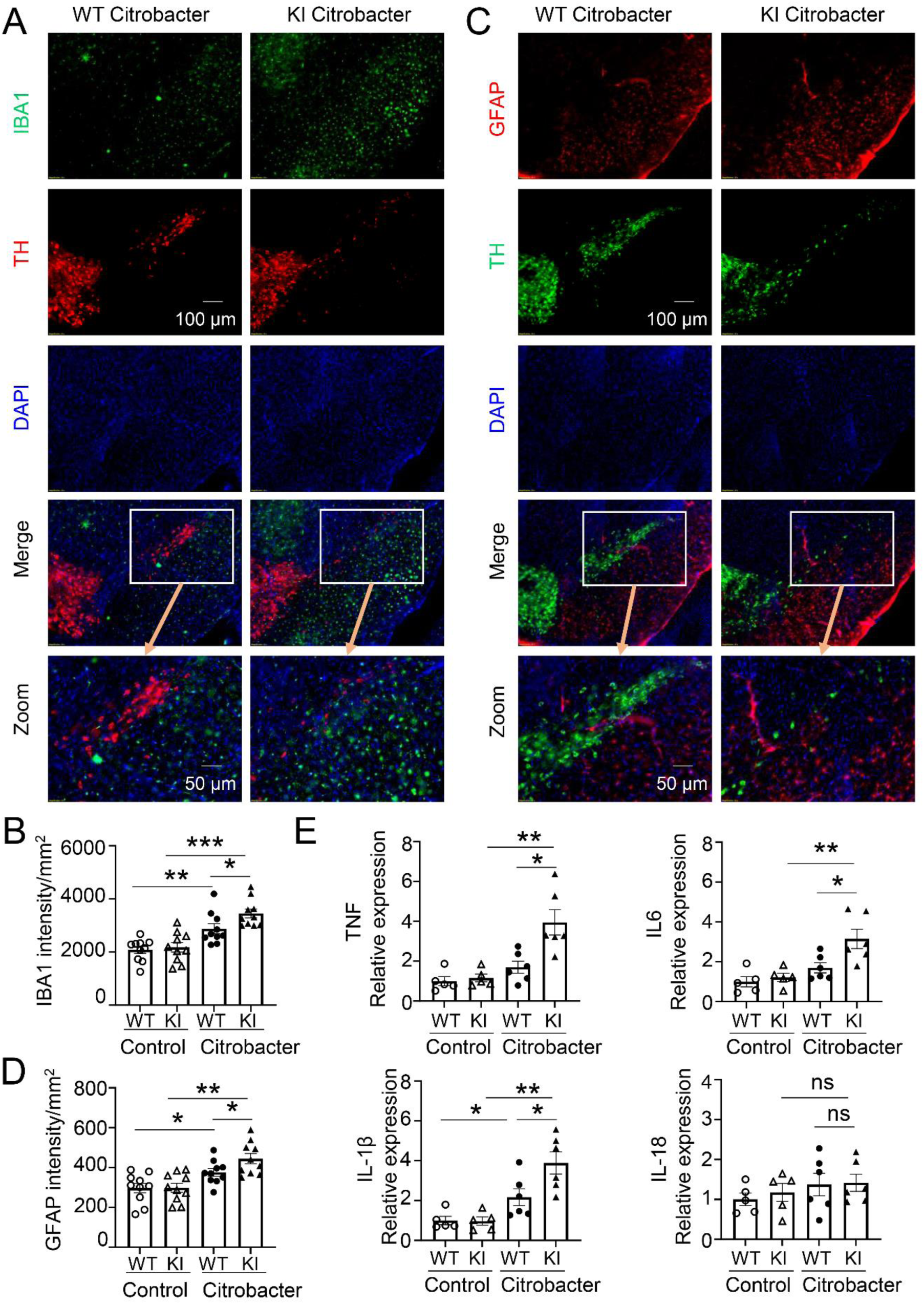
*C. rodentium* infection exacerbates neuroinflammation in LRRK2 G2019S KI mice. Neuroinflammatory responses were assessed in LRRK2 WT and LRRK2 G2019S KI mice at 6 months post-infection (mpi). Age- and genotype-matched uninfected littermates served as controls. (A) Representative immunofluorescence images of IBA1-positive microglia and TH in the substantia nigra (SN), with DAPI nuclear staining. Merged and higher-magnification views are shown. Scale bars, 100 μm (top) and 50 μm (zoom). (B) Quantification of IBA1 immunoreactivity in the SN, expressed as fluorescence intensity per mm^2^. (C) Representative immunofluorescence images of GFAP-positive astrocytes and TH in the SN, with DAPI staining. Merged and magnified views are shown. Scale bars, 100 μm (top) and 50 μm (zoom). (D) Quantification of GFAP immunoreactivity in the SN, expressed as fluorescence intensity per mm^2^. (E) qPCR analysis of pro-inflammatory cytokine expression in the midbrain dopaminergic regions (TNF, IL-6, IL-1β and IL-18). For (B) and (D), n = 10 mice per group; for (E), the control group n = 5 mice per group, the Citrobacter group n = 6 mice per group. Data are presented as mean ± SEM. *p<0.05, **p<0.01 and ***p<0.001. Data are representative of three independent experiments.

To further characterize inflammatory signaling, we quantified the expression of pro-inflammatory cytokines in the midbrain. Quantitative PCR analysis revealed robust upregulation of multiple inflammatory mediators in infected KI mice, including TNF-α, IL-6, and IL-1β, but not IL-18, compared with uninfected KI controls (**Fig. 3E**). Importantly, the induction of these cytokines was significantly greater in infected KI mice than in WT mice.

Collectively, these findings demonstrate that recurrent intestinal infection induces a robust neuroinflammatory response that is markedly exacerbated in LRRK2 G2019S KI mice. These results support a gene–environment interaction in which intestinal infection not only triggers neuroinflammation but also amplifies it in the context of pathogenic LRRK2 signaling, thereby generating a pro-inflammatory milieu that may contribute to subsequent dopaminergic neurodegeneration.

### Intestinal infection induces pathological α-synuclein accumulation in LRRK2 KI mice

Pathological aggregation of α-synuclein is a defining molecular hallmark of PD and is closely linked to dopaminergic neurodegeneration. Given that infected LRRK2 KI mice exhibited nigrostriatal degeneration following *C. rodentium* infection, we next examined whether this phenotype was accompanied by aberrant α-synuclein pathology in the midbrain. To this end, we assessed phosphorylated α-syn at serine 129 (p-αSyn), a well-established marker of pathogenic aggregation.

Strikingly, prominent p-αSyn–positive puncta and aggregates were predominantly observed in infected KI mice, whereas infected WT mice displayed only minimal background staining (**Fig. 4A**). These aggregates were largely localized to regions adjacent to, but not overlapping with, TH-positive neuronal cell bodies, suggesting preferential accumulation outside the SNpc somatic compartment. Quantitative analysis confirmed a significant increase in p-αSyn signal intensity in infected KI mice compared with both infected WT mice and uninfected KI controls (**Fig. 4B**).

**Figure 4.**
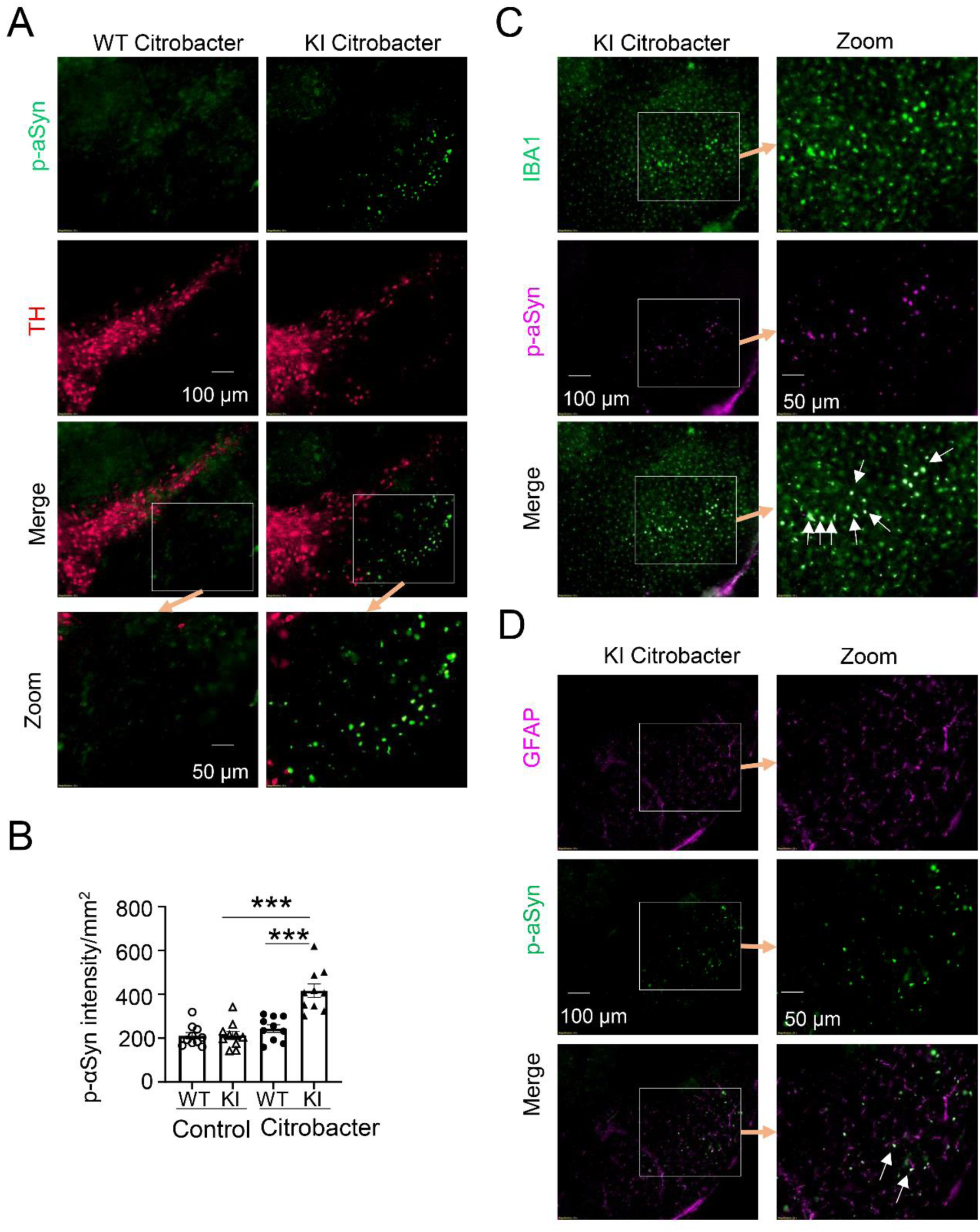
LRRK2 G2019S KI mice develop α-synuclein pathology in the substantia nigra following *C. rodentium* infection. Phosphorylated Ser129 α-synuclein (p-αSyn) was assessed in LRRK2 WT and LRRK2 G2019S KI mice at 6 months post-infection (mpi). Age- and genotype-matched uninfected littermates served as controls. (A) Representative immunofluorescence images of p-αSyn and TH staining in the substantia nigra (SN). Merged and higher-magnification views are shown. Scale bars, 100 μm (top) and 50 μm (zoom). (B) Quantification of p-αSyn immunoreactivity in the SN, expressed as fluorescence intensity per mm^2^. (C) Representative immunofluorescence images showing co-staining of p-αSyn with IBA1 (microglia) in the SN of KI mice following *C. rodentium* infection. Insets and higher-magnification views highlight regions of interest. Scale bars, 100 μm (left) and 50 μm (right). (D) Representative immunofluorescence images showing co-staining of p-αSyn with GFAP (astrocytes) in the SN of KI mice following infection. Insets and higher-magnification views are shown. Scale bars, 100 μm (left) and 50 μm (right). For (B), n = 10 mice per group. Data are presented as mean ± SEM***p<0.001. Data are representative of three independent experiments.

To further define the cellular context of α-syn pathology, we examined its spatial relationship with glial activation. p-αSyn–enriched regions in infected KI mice were closely associated with activated IBA1⁺ microglia (**Fig. 4C**) and, to a lesser extent, GFAP⁺ astrocytes (**Fig. 4D**), indicating a localized neuroinflammatory environment. Notably, p-αSyn signals were frequently observed in proximity to, but not extensively co-localized with, glial markers, suggesting that aggregates reside within inflamed microenvironments rather than exclusively within glial cells.

Collectively, these findings demonstrate that recurrent intestinal infection induces pathological α- syn accumulation in LRRK2 G2019S KI mice. The preferential localization of p-αSyn outside dopaminergic neuronal soma, together with its close association with activated glia, suggests that α-syn pathology emerges within localized neuroinflammatory microenvironments.

### Intestinal α-synuclein aggregation potentially links gut infection to brain pathology in LRRK2 KI mice

We hypothesized that the PD-like pathology observed in the brains of LRRK2 G2019S KI mice is associated with enhanced intestinal pathology. To test this, we examined whether intestinal α-syn pathology is altered under these conditions. We assessed p-αSyn accumulation in the colon. Immunofluorescence analysis revealed that recurrent *C. rodentium* infection induced a significant increase in p-αSyn signal in both wild-type (WT) and KI mice compared with their respective controls (**Fig. 5A, B**). Notably, this response was markedly amplified in KI mice, which exhibited substantially higher levels of p-αSyn deposition than infected WT animals. In WT mice, p-αSyn immunoreactivity was largely confined to the epithelial layer. In contrast, KI mice displayed pronounced expansion of p-αSyn signal beyond the epithelium into submucosal compartments (**Fig. 5A**), indicating a broader distribution of α-synuclein pathology. Collectively, these findings demonstrate that recurrent intestinal infection induces robust α-syn accumulation in the gut and markedly enhances both its magnitude and spatial distribution in LRRK2 G2019S KI mice.

**Figure 5.**
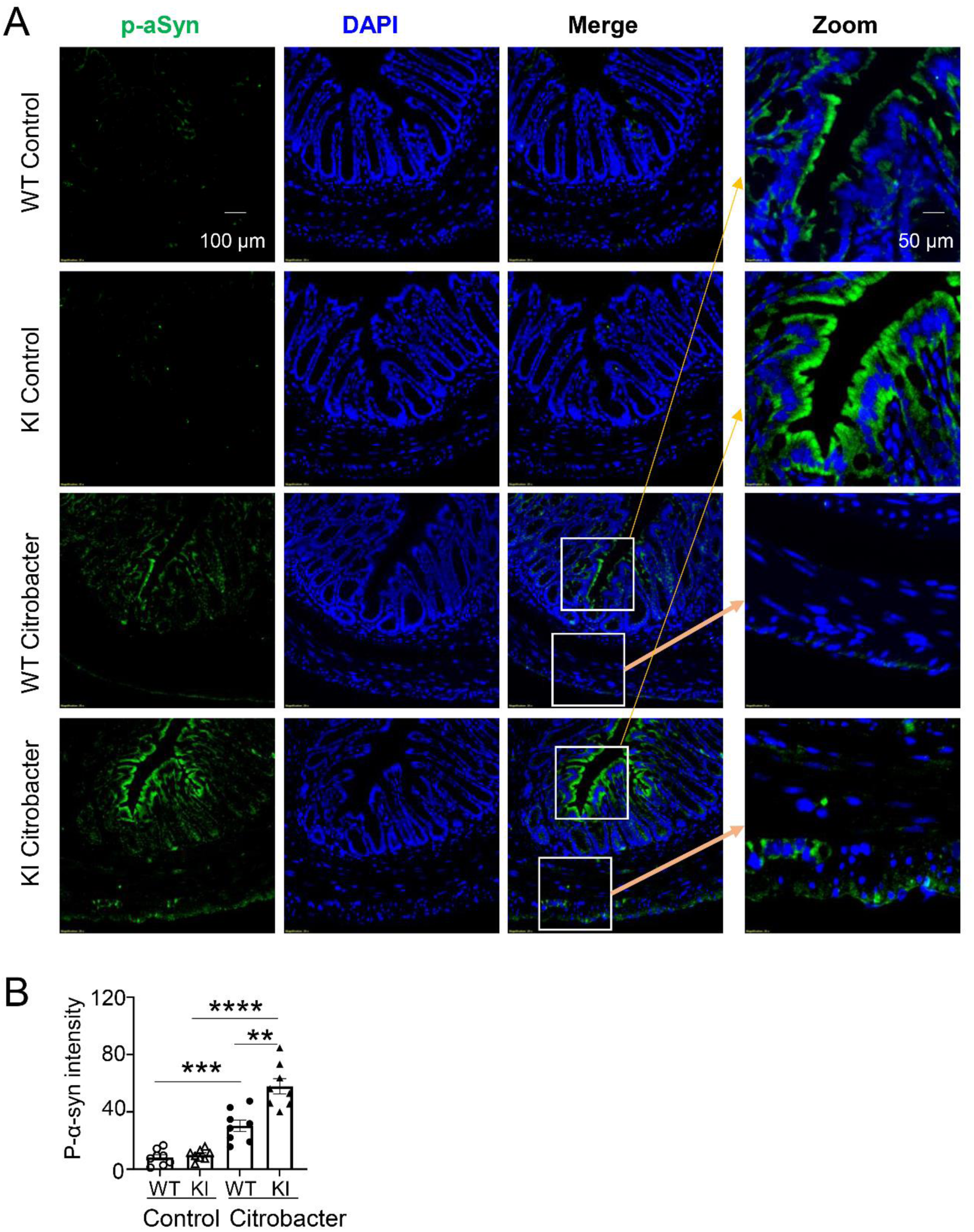
Enhanced gut α-synuclein pathology in LRRK2 G2019S KI mice following *C. rodentium* infection. p- áSyn was assessed in colon tissue from LRRK2 WT and LRRK2 G2019S KI mice at 6 months post-infection (mpi). Age- and genotype-matched uninfected littermates served as controls. (A) Representative immunofluorescence images of p-αSyn (green) and DAPI nuclear staining (blue) in colon sections. Merged and higher-magnification views highlight regions of interest. Scale bars, 100 μm (main images) and 50 μm (zoom). (B) Quantification of p-αSyn immunoreactivity in colon tissue, expressed as mean fluorescence intensity per unit area (n = 8 mice per group). Data are presented as mean ± SEM. **p<0.01,***p<0.001 and ****p<0.0001. Data are representative of three independent experiments.

### LRRK2 G2019S mutation exacerbates intestinal inflammation and barrier dysfunction following *C. rodentium* infection

Because repeated enteric infection induced pronounced intestinal α-synuclein accumulation in LRRK2 KI mice, we hypothesized that the pathogenic LRRK2 mutation increases susceptibility to intestinal inflammation during bacterial challenge. To test this hypothesis, WT and LRRK2 KI mice were subjected to a single cycle of *Citrobacter rodentium* infection and monitored longitudinally for pathological manifestations of colitis (**Fig. 6A**).

**Figure 6.**
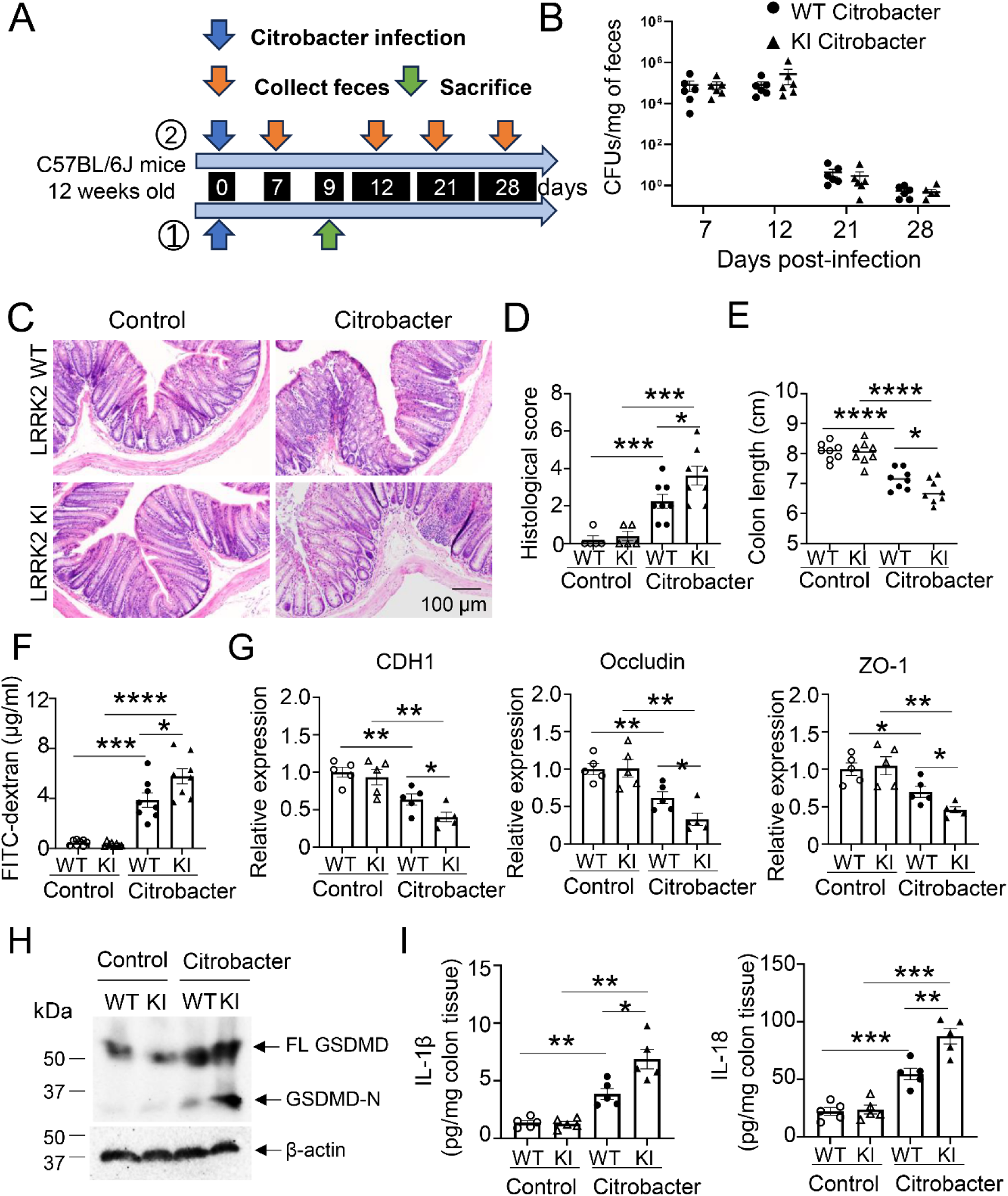
LRRK2 G2019S KI mice exhibit exacerbated gut inflammation following acute *C. rodentium* infection. (A) Experimental design. Twelve-week-old LRRK2 WT and LRRK2 G2019S KI mice were orally gavaged with C. rodentium (4 × 10^8^ CFU) on day 0. Fecal samples were collected at the indicated time points. One cohort of mice was sacrificed on day 9 post-infection for tissue analyses, whereas a separate cohort was followed longitudinally for fecal sample collection through day 28 post-infection. (B) Fecal bacterial burden (CFU per mg feces) measured at days 7, 12, 21, and 28 post- infection. (C) Representative H&E-stained sections of colon tissue. Scale bar, 100 μm. (D) Histopathological scoring of colonic tissue. (E) Colon length measurements. (F) Intestinal permeability assay. Mice were gavaged with FITC-dextran, and serum FITC- dextran levels were measured 4 h later to assess gut barrier integrity. (G) qPCR analysis of tight junction–associated genes (CDH1, Occludin, and ZO-1) in colon tissue. (H) Western blot analysis of gasdermin D (GSDMD) cleavage in colon explant tissue, with β- actin as an internal control. (I) Quantification of IL-1β and IL-18 levels in colon explant tissues by ELISA, expressed as pg/mg colon tissue. For (B), n = 10 mice per group; For (D) control groups n = 5 mice, Citrobacter groups n=8 mice; For (E–F), n = 8 mice per group; for (G) and (I), n = 5 mice per group. Data are presented as mean ± SEM, *p<0.05, **p<0.01 and ***p<0.001. Data are representative of three independent experiments.

Quantification of fecal bacterial burden demonstrated a typical course of *C. rodentium* infection, with colony-forming units (CFUs) peaking at day 7 post-infection and progressively declining thereafter, consistent with normal bacterial clearance kinetics. Importantly, no significant differences in fecal bacterial loads were observed between infected WT and KI mice at any examined time point (**Fig. 6B**), indicating that the LRRK2 G2019S mutation does not impair host bacterial clearance.

Despite comparable pathogen burdens, infected KI mice developed markedly more severe intestinal pathology than infected WT controls. Histopathological analysis of colonic tissues collected at day 9 post-infection revealed substantially exacerbated colitis in KI mice, characterized by extensive epithelial erosion, crypt architectural destruction, mucosal ulceration, and dense inflammatory cell infiltration within the lamina propria (**Fig. 6C**). Semi-quantitative histological scoring further confirmed significantly increased inflammatory severity in infected KI mice relative to infected WT animals (**Fig. 6D**). Consistent with enhanced intestinal inflammation, infected KI mice also exhibited significantly shortened colons compared with infected WT mice (**Fig. 6E**), a well-established indicator of colonic inflammatory injury.

Given that epithelial barrier dysfunction is a critical determinant of intestinal inflammation and gut– brain communication, we next assessed intestinal permeability using FITC–dextran assays. Following oral administration of FITC–dextran, infected KI mice exhibited significantly elevated serum FITC–dextran levels compared with infected WT controls (**Fig. 6F**), indicating increased intestinal permeability and impaired epithelial barrier integrity.

To further define the molecular basis underlying barrier disruption, we examined the expression of epithelial junctional genes in colonic tissues. Quantitative PCR analysis demonstrated significant downregulation of multiple epithelial barrier-associated genes, including *Cdh1, Occludin, and ZO-1*, in infected KI mice relative to infected WT controls (**Fig. 6G**), supporting the presence of compromised tight junction integrity and epithelial barrier disruption.

Because inflammasome-mediated inflammatory signaling has been implicated in both intestinal pathology and neuroinflammatory processes, we next evaluated inflammasome activation in the colon upon *C. rodentium* infection. Immunoblot analysis revealed enhanced cleavage of gasdermin D (GSDMD), including accumulation of the active N-terminal fragment (GSDMD-N), in infected KI mice compared with WT animals (**Fig. 6H**), indicating increased pyroptotic signaling. Consistently, ELISA analysis demonstrated significantly elevated production of IL-1β and IL-18 in colonic tissues from infected KI mice (**Fig. 6I**), further supporting enhanced inflammasome activation in the setting of pathogenic LRRK2 signaling.

Collectively, these findings demonstrate that the LRRK2 G2019S mutation sensitizes the intestinal mucosa to exaggerated inflammatory injury and epithelial barrier disruption following enteric bacterial infection. Notably, these pathological alterations occurred despite comparable bacterial clearance between WT and KI mice, suggesting that mutant LRRK2 primarily amplifies host inflammatory responses rather than impairing antimicrobial defense. This heightened inflammatory and hyperpermeable intestinal environment may contribute to the enhanced intestinal α-syn accumulation observed in KI mice, thereby facilitating pathogenic gut–brain axis signaling associated with PD-related neuroinflammation.

## Discussion

In the present study, we establish a gene–environment interaction model demonstrating that recurrent enteric bacterial infection can trigger progressive Parkinson’s disease (PD)-like pathology in genetically susceptible **LRRK2 G2019S** knock-in (KI) mice. Repeated *Citrobacter rodentium* infection induced progressive motor impairment, selective nigrostriatal dopaminergic neurodegeneration, neuroinflammation, and pathological α-synuclein accumulation in **LRRK2 G2019S** KI mice, whereas wild-type animals remained comparatively resistant to these pathological changes. These observations are consistent with the pioneering study by Matheoud et al. demonstrating that recurrent intestinal bacterial infection can trigger PD-like motor and neuropathological abnormalities in genetically susceptible **Pink1**^−/−^ mice (25), supporting the concept that enteric infection can interact with PD-associated genetic susceptibility to promote neurodegeneration. Our results extend this gene–environment paradigm to the **LRRK2 G2019S** mutation, indicating that distinct PD-associated genetic risk factors can converge with chronic enteric infection to drive disease pathogenesis. Collectively, these findings suggest that pathogenic LRRK2 signaling acts as a sensitizing factor that requires sustained inflammatory challenge to promote disease progression. Mechanistically, infected KI mice developed exaggerated intestinal inflammation, epithelial barrier dysfunction, and inflammasome activation despite normal bacterial clearance, further supporting a gut–brain axis mechanism linking chronic intestinal inflammation to PD-related neurodegeneration.

Importantly, the recurrent infection paradigm used in this study may more closely reflect the chronic and intermittent intestinal inflammatory exposures experienced throughout human aging and environmental pathogen encounters. Unlike acute inflammatory models, repeated mucosal infection may progressively establish long-term inflammatory priming, sustain innate immune activation, and cumulative tissue injury that gradually overcome compensatory neuroimmune resilience mechanisms. In genetically susceptible hosts, such chronic inflammatory sensitization may create a permissive environment for persistent α-synuclein pathology, neuroinflammation, and progressive neurodegeneration. These findings support the emerging concept that repeated peripheral inflammatory insults, rather than a single acute event, may represent an important environmental driver of PD progression(21).

Previous studies demonstrated that intracerebral inoculation of α-synuclein preformed fibrils (PFFs) in wild-type mice is sufficient to induce progressive PD-like pathology, including pathological α-synuclein propagation, p-αSyn accumulation, nigrostriatal dopaminergic neurodegeneration, and motor dysfunction, supporting a central role for α-synuclein-mediated neurotoxicity in PD pathogenesis(27). Importantly, pathogenic LRRK2 G2019S signaling further exacerbates PFF-induced α-synuclein aggregation, p-αSyn pathology, and neurodegeneration, suggesting that LRRK2 enhances susceptibility to α-synuclein-associated neurotoxicity(28, 29). Consistent with these findings, recurrent enteric infection in our model induced robust endogenous p-αSyn accumulation in the brains of LRRK2 G2019S KI mice. This pathological α- synuclein accumulation may contribute to the progressive PD-like phenotypes observed in this model. Interestingly, and distinct from classical neuron-centered α-synucleinopathy models, p- αSyn aggregates in our model preferentially localized to inflammatory niches adjacent to activated microglia and astrocytes rather than directly overlapping with dopaminergic neuronal soma. This unique distribution pattern suggests that chronic neuroinflammatory microenvironments may play an important role in promoting α-synuclein phosphorylation, aggregation, and/or impaired clearance during early disease progression.

Compared with previously reported gut–brain axis LRRK2-associated PD models, our model demonstrates several important advantages. In the Dextran sodium sulfate (DSS)-induced chronic colitis model using LRRK2 G2019S transgenic mice, chronic intestinal inflammation induced microglial activation, dopaminergic neuronal loss, and locomotor dysfunction through TNF-α-dependent inflammatory pathways, supporting an important role for intestinal inflammation in PD-related neurodegeneration(20). However, despite increased total colonic α-synuclein expression, that study failed to detect phospho-Ser129 α-synuclein inclusions or Lewy body-like pathology in brain tissues(20), suggesting that the DSS model primarily reflects inflammation-driven neurodegeneration without robust endogenous synucleinopathy. Another related study using LRRK2 G2019S KI mice demonstrated that chronic DSS colitis could further exacerbate neurodegeneration in the presence of direct intracranial α-synuclein overexpression induced by p-αSyn/AAV α-syn administration(21). While these findings support an interaction between intestinal inflammation and α-synuclein-mediated neurotoxicity, PD-like pathology in that model still depended on artificial induction of brain α-synuclein burden. In contrast, our model demonstrates that recurrent enteric infection alone is sufficient to induce robust endogenous p- αSyn accumulation in both intestinal and brain tissues together with chronic neuroinflammation and progressive dopaminergic neurodegeneration, without requiring exogenous α-synuclein overexpression or fibril inoculation.

Furthermore, compared with the previously reported *E. coli*-induced PD model in LRRK2 mutant mice, in which p-αSyn accumulation was largely restricted to the intestinal epithelial layer(30), recurrent *C. rodentium* infection in our model induced extensive endogenous p-αSyn accumulation extending beyond the epithelium into submucosal compartments, suggesting progressive involvement of deeper intestinal neuroimmune structures. Since the intestinal submucosa contains dense enteric neuronal, immune, glial, and vascular networks, such pathology may facilitate inflammatory amplification and pathological gut-to-brain propagation. Collectively, these findings establish our model as a physiologically relevant gut–brain axis PD model that more closely recapitulates key pathological features of progressive PD, including endogenous α-synuclein pathology, chronic neuroinflammation, and gut-to-brain neurodegenerative progression.

The enhanced inflammasome activation observed in infected LRRK2 G2019S KI mice suggests a potential mechanistic link between pathogenic LRRK2 signaling, chronic intestinal inflammation, and PD-related neurodegeneration. Recent studies demonstrated that LRRK2 G2019S enhances neutrophil recruitment, Neutrophil Extracellular Trap (NET) formation, and Th17-associated inflammatory responses during *C. rodentium* infection (31), supporting the concept that pathogenic LRRK2 signaling broadly sensitizes innate immune pathways to enteric inflammatory stimuli. Our findings further demonstrated increased GSDMD cleavage together with elevated IL- 1β and IL-18 production in colonic tissues, indicating enhanced pyroptotic and inflammasome-mediated inflammatory signaling. Chronic inflammasome activation may contribute to disease progression through multiple converging mechanisms, including cytokine-driven systemic inflammation, epithelial barrier disruption, immune cell recruitment, and amplification of α- synuclein pathology(32). Increased intestinal permeability may further permit translocation of microbial products, inflammatory mediators, and potentially pathogenic protein aggregates into systemic circulation and neural pathways, thereby sustaining chronic peripheral and central immune activation(33). Together, these findings support a model in which pathogenic LRRK2 signaling amplifies inflammasome activation and barrier dysfunction, facilitating gut-derived inflammatory signaling that contributes to progressive neurodegeneration.

Several limitations of this study should also be acknowledged. Although our data strongly support a gut–brain axis mechanism, the precise pathways linking intestinal inflammation to central neurodegeneration remain to be fully defined. Future studies will be required to determine whether vagal transmission(34), circulating inflammatory mediators, microbial metabolites, or direct propagation of α-synuclein contribute to disease progression in this model. In addition, while p- αSyn accumulation and neurodegeneration were observed following recurrent infection, the temporal sequence linking intestinal pathology to brain pathology warrants further mechanistic investigation. Finally, whether pharmacological inhibition of LRRK2 kinase activity or inflammasome signaling can prevent infection-induced pathology remains an important therapeutic question.

In summary, our study establishes recurrent enteric bacterial infection as an environmental trigger that cooperates with pathogenic LRRK2 signaling to drive progressive PD-like pathology. These findings support a unifying model in which chronic intestinal inflammatory insults synergize with PD-associated genetic susceptibility to promote α-synuclein accumulation, neuroinflammation, and nigrostriatal neurodegeneration through gut–brain immune mechanisms. Collectively, our data establish this model as a physiologically relevant and mechanistically informative platform for studying PD pathogenesis and gut–brain axis-mediated neurodegeneration.

## Abbreviations

### Abbreviation Full Term

PD: Parkinson’s disease
α-syn / αSyn: Alpha-synuclein
p-αSyn: Phosphorylated alpha-synuclein
pS129: Phosphorylated serine 129
LRRK2: Leucine-rich repeat kinase 2
KI: Knock-in
WT: Wild type
SNpc: Substantia nigra pars compacta
SNpr: Substantia nigra pars reticulata
VTA: Ventral tegmental area
TH: Tyrosine hydroxylase
GFAP: Glial fibrillary acidic protein
IBA1: Ionized calcium-binding adapter molecule 1
DSS: Dextran sodium sulfate
PFFs: Preformed fibrils
IBD: Inflammatory bowel disease
CFU / CFUs: Colony-forming unit(s)
Mpi: Months post-infection
p.i.: Post-infection
PBS: Phosphate-buffered saline
PFA: Paraformaldehyde
OCT: Optimal cutting temperature
FITC-dextran: Fluorescein isothiocyanate-dextran
OD600: Optical density at 600 nm
LB: Luria-Bertani
DMEM: Dulbecco’s Modified Eagle Medium
RPMI: Roswell Park Memorial Institute medium
FBS: Fetal bovine serum
DAPI: 4′,6-diamidino-2-phenylindole
ROI / ROIs: Region(s) of interest
H&E: Hematoxylin and eosin
ELISA: Enzyme-linked immunosorbent assay
RIPA: Radioimmunoprecipitation assay
BCA: Bicinchoninic acid
SDS-PAGE: Sodium dodecyl sulfate-polyacrylamide gel electrophoresis
PVDF: Polyvinylidene fluoride
HRP: Horseradish peroxidase
RNA: Ribonucleic acid
cDNA: Complementary DNA
Q-PCR: Quantitative polymerase chain reaction
TNF-α: Tumor necrosis factor alpha
IL: Interleukin
GSDMD: Gasdermin D
GSDMD-N: N-terminal fragment of gasdermin D
Cdh1: Cadherin 1
ZO-1: Zonula occludens-1

## Acknowledgments

We sincerely thank the Comparative Pathology Laboratory (CPL) and Histology Research Laboratory (HPL) in the Department of Pathology at the University of Iowa for providing outstanding necropsy and histology. We sincerely thank Shane Heiney from the Neural Circuits and Behavior Core at the University of Iowa for his assistance with grip strength testing and open field test analysis. We sincerely thank Kartik Sivakumar for technical assistance with microscopy setup, cryostat sectioning, and immunostaining of brain tissue samples.

## Authors’ Contributions

Y.W. designed and performed the experiments, prepared the figures, interpreted the data, and wrote the manuscript. M.W. contributed to the pathology analysis, data interpretation and manuscript revision. N.C., L.Z, S.L., M.B., T.B., M.J. and T.S. assisted in mouse genotyping, immunoblotting, and real-time PCR and behavior testing. R.T. and W.W. contributed to the pathology analysis. N.N. contributed to project coordination, data interpretation, and manuscript revision. Z.K. was integral for experimental design, manuscript writing, data interpretation, and project coordination.

## Data availability

All data are available upon reasonable requests made to the corresponding author.

## Ethics approval and consent to participate

Animal experiments were approved by the Institutional Animal Care and Use Committee (IACUC) of the University of Iowa.

