## Supplemental Table 1 and 2 for "Gut bacterial Infection drives Parkinsonian pathology in LRRK2 G2019S Knock-in Mice" (from another version of this preprint: version 1, 2026-06-10)

Table 1. First antibodies used for IHC

| **Host** | **Antibodies** | **Catalog number; Company** |
| --- | --- | --- |
| Rabbit | anti-tyrosine hydroxylase (TH) antibody | AB152, Sigma-Aldrich |
| Chicken | anti-tyrosine hydroxylase (TH) antibody | ab764422, Abcam |
| Rabbit | anti–phosphorylated α-syn (p-α-syn, Ser129) | ep1536y, AB51253, Abcam |
| Mouse | Purified anti-α-Synuclein Phospho (Ser129) Antibody | 825701; BioLegend |
| Rabbit | anti-Ionized calcium-binding adapter molecule 1 (IBA1) antibody | 019-19741,FUJIFILM Wako Pure Chemical Corporation) |
| Mouse | anti–glial fibrillary acidic protein (GFAP) antibody | MAB3402; Sigma-Aldrich |

Table 2. Second antibodies used for IHC

|  | **Antibodies** | **Catalog number; Company** |
| --- | --- | --- |
| Anti-Rabbit | Donkey anti-Rabbit IgG (H+L) Highly Cross-Adsorbed Secondary Antibody, Alexa Fluor™ 488 | A21206; Invitrogen |
|  | Goat anti-Rabbit IgG (H+L) Cross-Adsorbed Secondary Antibody, Alexa Fluor™ 594 | A11012; Invitrogen |
|  | Goat anti-Rabbit IgG (H+L) Cross-Adsorbed Secondary Antibody, Alexa Fluor™ 680 | A21076; Invitrogen |
| Anti-Mouse | Goat anti-Mouse IgG (H+L) Highly Cross-Adsorbed Secondary Antibody, Alexa Fluor™ Plus 488 | A32723TR; Invitrogen |
|  | Anti-mouse IgG (H+L), F(ab')2 Fragment (Alexa Fluor® 594 Conjugate) | 8890S; CST |
|  | Goat anti-Mouse IgG (H+L) Cross-Adsorbed Secondary Antibody, Alexa Fluor™ 680 | A21057; Invitrogen |
| Anti-Rat | Donkey anti-Rat IgG (H+L) Highly Cross-Adsorbed Secondary Antibody, Alexa Fluor™ 594 | A21209; Invitrogen |
| Anti-chicken | Goat anti-Chicken IgY (H+L) Cross-Adsorbed Secondary Antibody, DyLight™ 594 | SA5-10072; Invitrogen |
